# Acetylcholine boosts dendritic NMDA spikes in a CA3 pyramidal neuron model

**DOI:** 10.1101/2021.03.01.433406

**Authors:** Rachel Humphries, Jack R. Mellor, Cian O’Donnell

## Abstract

Acetylcholine has been proposed to facilitate the formation of memory ensembles within the hippocampal CA3 network, by enhancing plasticity at CA3-CA3 recurrent synapses. Regenerative NMDA receptor (NMDAR) activation in CA3 neuron dendrites (NMDA spikes) increase synaptic Ca^2+^ influx and can trigger this synaptic plasticity. Acetylcholine inhibits potassium channels which enhances dendritic excitability and therefore could facilitate NMDA spike generation. Here, we investigate NMDAR-mediated nonlinear synaptic integration in stratum radiatum (SR) and stratum lacunosum moleculare (SLM) dendrites in a reconstructed CA3 neuron computational model and study the effect of acetylcholine on this nonlinearity. We found that distal SLM dendrites, with a higher input resistance, had a lower threshold for NMDA spike generation compared to SR dendrites. Simulating acetylcholine by blocking potassium channels (M-type, A-type, Ca^2+^-activated, and inwardly-rectifying) increased dendritic excitability and reduced the number of synapses required to generate NMDA spikes, particularly in the SR dendrites. The magnitude of this effect was heterogeneous across different dendritic branches within the same neuron. These results predict that acetylcholine facilitates dendritic integration and NMDA spike generation in selected CA3 dendrites which could strengthen connections between specific CA3 neurons to form memory ensembles.

**Highlights:**

- Using biophysical computational models of CA3 pyramidal neurons we estimated the quantitative effects of acetylcholine on nonlinear synaptic integration.
- Nonlinear NMDA spikes can be triggered by fewer synapses in distal dendrites due to increased local input resistance.
- Acetylcholine broadly reduces the number of synapses needed to trigger NMDA spikes, but the magnitude of the effect varies across dendrite branches within a single neuron.
- No single potassium channel type is the dominant mediator of the excitability effects of acetylcholine.

## Introduction

Episodic memories are encoded in the hippocampus by processing information from the entorhinal cortex within the dentate gyrus, CA3, CA2 and CA1 regions. Each region performs a distinct role for memory processing (Marr, 1971; McClelland and Goddard, 1996; Kesner and Rolls, 2015). CA3 is unique within the hippocampus because its excitatory pyramidal neurons are recurrently connected, allowing the emergence of CA3 networks with attractor dynamics (Marr, 1971; Hopfield, 1982; Kesner and Rolls, 2015; Guzman et al., 2016). Computationally, such networks are important for memory retrieval since they allow sparse external cues to drive attractor networks towards stable memory states; a process known as pattern completion (Gold and Kesner, 2005; Yassa and Stark, 2011). In order for the network to form new memory associations, it needs to adapt which neurons participate by selectively strengthening or weakening the recurrent excitatory synapses between CA3 pyramidal cells (Treves and Rolls, 1994; Tsodyks, 1999; Nakazawa et al., 2002; Rebola et al., 2017). In these cells, synaptic plasticity can be triggered by the coincident firing of action potentials (Debanne et al., 1998; Mishra et al., 2016), but recent evidence also suggests that dendritic spikes of Ca^2+^ caused by nonlinear synaptic integration and NMDA receptor (NMDAR) activation are highly effective at inducing synaptic plasticity (Golding et al., 2002; Brandalise et al., 2016; Mishra et al., 2016; Weber et al., 2016).

There are 3 major excitatory inputs to CA3 pyramidal neurons: mossy fiber inputs from dentate gyrus granule cells, perforant path inputs from entorhinal cortex layer II and recurrent associational/commissural inputs from CA3 pyramidal neurons (Witter, 2007; Hunt et al., 2018). Each of these inputs synapse onto different regions of the dendritic tree: mossy fibers onto the most proximal dendrites in the stratum lucidum (SL); perforant path onto the most distal dendrites in the stratum lacunosum-moleculare (SLM); and associational/commissural onto the mid region of the apical dendrites in the stratum radiatum (SR) as well as onto the basal dendrites in the stratum oriens (SO). Nonlinear synaptic summation and dendritic Ca^2+^ spikes can arise from near coincident activity at synapses either within or between these pathways (Makara and Magee, 2013; Brandalise and Gerber, 2014; Brandalise et al., 2016; Weber et al., 2016) but the precise configurations of synaptic activity required to drive dendritic spikes have not been fully explored.

Acetylcholine is released in response to salient, rewarding or arousing stimuli that require learning of new associations (Hasselmo, 2006; Teles-Grilo Ruivo and Mellor, 2013; Lovett-Barron et al., 2014; Hangya et al., 2015; Teles-Grilo Ruivo et al., 2017) and therefore is predicted to facilitate the formation of new memory ensembles in the hippocampus (Hasselmo et al., 1996; Hasselmo, 1999; Meeter et al., 2004; Prince et al., 2016). In line with this, acetylcholine facilitates NMDAR function and induction of synaptic plasticity (Markram and Segal, 1992; Marino et al., 1998; Buchanan et al., 2010; Fernández De Sevilla and Buño, 2010; Gu and Yakel, 2011; Dennis et al., 2016; Papouin et al., 2017). Disruption of cholinergic signalling impairs memory encoding (Berger-Sweeney et al., 2001; Anagnostaras et al., 2003; Rogers and Kesner, 2003). Importantly, activation of muscarinic M1 receptors on CA3 pyramidal neurons depolarizes the membrane potential and increases input resistance (Sun and Kapur, 2012). This is expected to increase cell excitability and is observed for regular spiking pyramidal neurons, but a sub-population of CA3 pyramidal cells respond specifically by reducing burst firing (Hunt et al., 2018). These effects result from inhibition of potassium channels including: M-type (Km) (Delmas and Brown, 2005), A-type (Ka) (Hoffman and Johnston, 1998), G-protein-coupled inwardly-rectifying (GIRK, Kir) (Sohn et al., 2007) and small-conductance Ca^2+^-activated channels (Kca (SK)) (Buchanan et al., 2010; Giessel and Sabatini, 2010). Integration of synaptic inputs and the nonlinear generation of dendritic spikes are highly sensitive to the intrinsic excitability of dendrites (Gulledge et al., 2005; Stuart and Spruston, 2015). Dendritic integration properties vary in different regions of the pyramidal neuron because of the shape of the dendritic tree and its non-uniform distribution of voltage-gated ion channels (Spruston, 2008). Since neuromodulators such as acetylcholine regulate neuronal membrane conductances, they may also control dendritic spike generation and therefore synaptic plasticity and CA3 network ensemble formation (Prince et al., 2016; Fernández de Sevilla et al., 2020). Of the potassium channels that acetylcholine inhibits, Ka, Kca and Kir have been shown to modulate NMDAR-mediated dendritic spikes (NMDA spikes) (Cai et al., 2004; Makara and Magee, 2013; Malik and Johnston, 2017). Additionally, acetylcholine facilitates dendritic integration in neocortical layer 5b pyramidal neurons (Williams and Fletcher, 2019) and in hippocampal OLM interneurons (Hagger-Vaughan and Storm, 2019).

Given the above findings, it is possible that acetylcholine enables CA3 network plasticity by two distinct mechanisms: first, by directly depolarising dendrites to increase local NMDAR activation and dendritic spine Ca^2+^ influx; or second, by boosting neural spiking and inducing Hebbian plasticity between recurrently connected neurons. In either case, it will be crucial to understand how acetylcholine affects dendritic integration and NMDA spikes in CA3 pyramidal neurons. We addressed this problem using computational models of 2-compartment and detailed reconstructed CA3 pyramidal neurons, exploring dendritic integration of both perforant path and SR recurrent associational/commissural inputs. We found that the degree of synaptic integration depended on distance from the soma and that simulated acetylcholine enhanced dendritic NMDA spikes by reducing the number of synapses required to cause nonlinear summation.

## Methods

All simulations were performed using NEURON software (Carnevale and Hines, 2006) and Python 3.8.

### 2-compartment model

For the initial simulations, we used a 2-compartment model with a soma (500 μm^2^ area) and a dendrite (1 μm diameter x 200 μm length). We set the axial resistivity to 150 MΩ and the leak reversal potential to −65 mV. We took the channels included from a previous CA3 neuron model (Hemond et al., 2008) and included a sodium channel, several potassium channels (A-type, M-type, delayed-rectifier), Ca^2+^ channels (L-type, N-type, T-type), Ca^2+^-activated potassium channels (K_AHP_, BK) and a hyperpolarization-activated current (I_h_). For these simulations, we set the sodium channel conductance to zero to prevent action potentials. In order to capture the effects of acetylcholine, we added two extra conductances to this model: a small-conductance Ca^2+^-activated potassium channel (Kca or SK channel; (Combe et al., 2018)) and an inwardly-rectifying potassium channel (Kir or GIRK channel; (Martinello et al., 2019). To build the model with reasonable contributions of the relevant potassium channels, the conductance of each channel was optimised to best fit with experimental data (Figure 1). This was done by implementing a covariance matrix adaptation evolution strategy (Hansen, 2016; Jȩdrzejewski-Szmek et al., 2018) in which parameters were adjusted in order to match outputs from the model as closely as possible with experimental measurements of resting membrane potential and input resistance. The optimisation was implemented using the PyCMA package (https://github.com/CMA-ES/pycma). Here, we optimised the Ka, Km, Kca, Kir and leak peak conductances, as well as shifts in the voltage-activation curves for Ka, Km and Kir channels, using published data describing the changes in membrane potential and input resistance upon blocking each of the channels experimentally (Figure 1, (Sun and Kapur, 2012; Makara and Magee, 2013)). The experimental data was obtained from two different studies, with the Ka, Kca and Kir data from one study (Makara and Magee, 2013) and the Km and acetylcholine receptor data from a separate study (Sun and Kapur, 2012). To account for this, the target membrane potential and input resistance values for Km and acetylcholine were adjusted slightly to reflect the different resting membrane potentials of the two experimental set ups. Peak conductances for all channels are displayed in Table 1. NMDAR-containing synapses (NMDA synapses) were modelled with a dual exponential model that included a voltage-dependent magnesium block as in Baker et al., (2011). The NMDA synapse had a rise time of 4 ms and a decay time of 42 ms.

**Figure 1:**
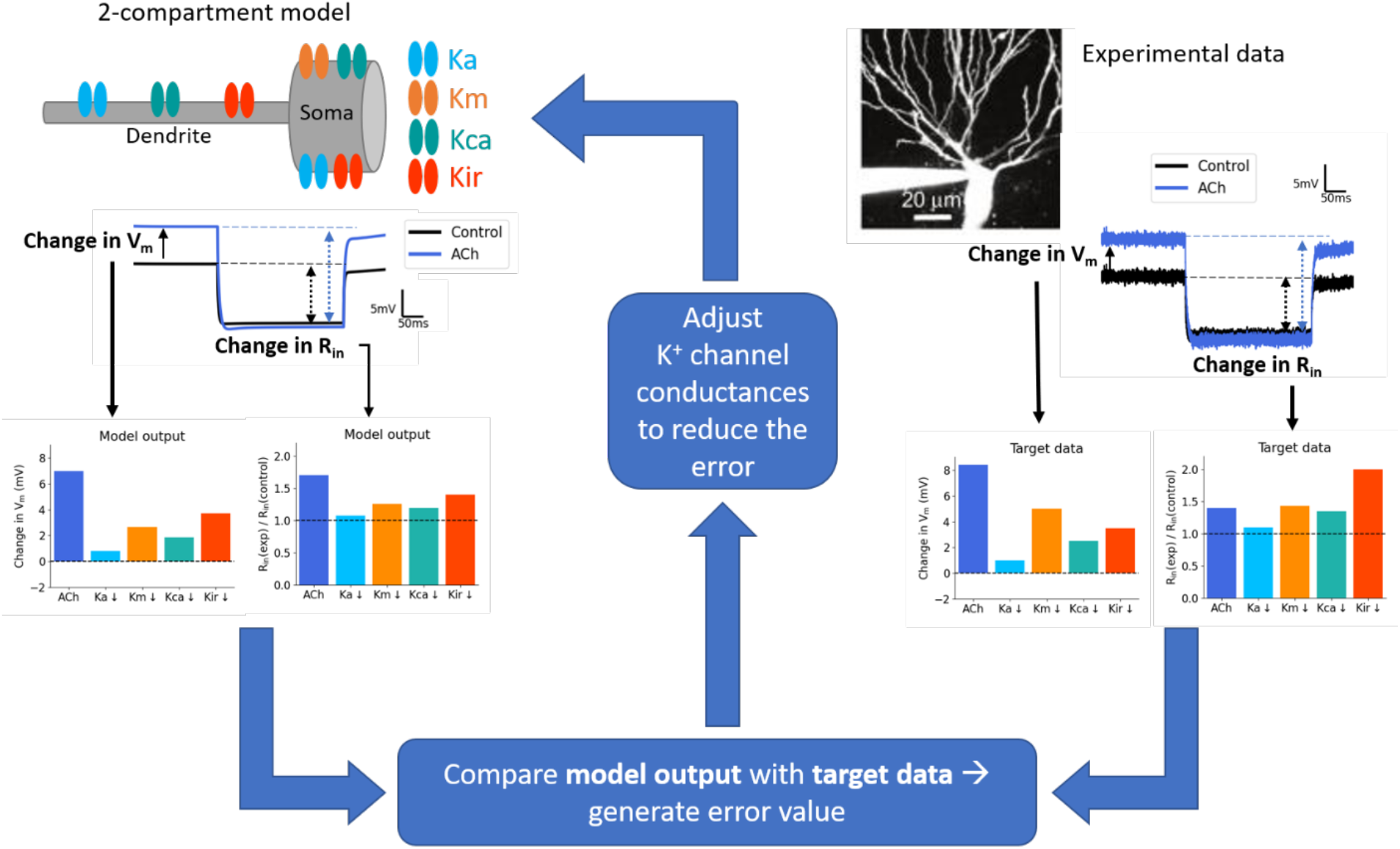
Optimisation of potassium conductances in 2-compartment model. Potassium channel conductance parameters were optimised by fitting to experimental data (from Makara and Magee (2013) and Sun and Kapur (2012)) measuring the change in membrane potential and input resistance after inhibiting the relevant potassium conductance. The error represented the difference between the model output and the target data which then fed back into the optimisation algorithm to adjust the potassium conductances. This cycle was repeated many times until the error was minimised. The bar plots shown represent the target experimental data used (right) and the final optimised output in the model (left). Neuron image adapted from Makara and Magee (2013). V_m_ = resting membrane potential; R_in_ = input resistance; Ka = A-type potassium channel, Km = M-type potassium channel, Kca = SK channel, Kir = GIRK channel.

**Table 1:** Peak conductance parameters for all channels. Ka, Km, Kca, Kir and the leak peak conductances were optimised for both models as shown in Figure 1. Kdr and Ih peak conductances were additionally optimised in the reconstructed neuron model. The remaining conductance values were taken from Hemond et al., (2008).

| Channel | Peak conductance (S/cm<sup>2</sup>) |  |
| --- | --- | --- |
|  | 2-compartment model | Reconstructed neuron model |
| Ka | 0.398 | $5.56 \times 10^{-11}$ |
| Km | 0.033 | 0.0011 |
| Kca | 0.035 | 0.0012 |
| Kir | 0.0012 | $3.47 \times 10^{-6}$ |
| Leak (gpas) | $7.25 \times 10^{-5}$ | $4.4 \times 10^{-10}$ |
| Kdr | 0.005 | $4.6 \times 10^{-9}$ |
| Na | 0.0 | 0.0 |
| CaT, CaN, CaL | $1 \times 10^{-5}$ | $1 \times 10^{-5}$ |
| Kc | $5 \times 10^{-5}$ | $5 \times 10^{-5}$ |
| Kahp | 0.0001 | 0.0001 |
| Ih | 0.00001 | $5.6 \times 10^{-6}$ |

### Reconstructed CA3 pyramidal neuron model

This model was based on the 943-compartment model from Hemond et al. (2008), reconstructed from a filled rat CA3 pyramidal neuron. It contained the same conductances as the 2-compartment model but the distribution of the Ka and Kir conductance varied along the dendritic arbour, as reported experimentally (Kim et al., 2012; Degro et al., 2015). The optimisation process of the potassium conductances (Figure 1) was repeated separately for this model (Table 1). To adjust for the lack of dendritic spines in the model, the membrane capacitance and resistance were doubled and halved, respectively, in all dendritic compartments. The dendritic regions were divided based on their distance from the soma (SL dendrites < 150μm; SR dendrites 150-400μm; SLM dendrites > 400μm, Figure 3A). AMPAR-containing synapses (AMPA synapses), with a rise and decay time of 0.5 and 1.5 ms, respectively, and 0.5x the peak conductance of NMDA synapses, were also included in this model (Table 2). Synapses were positioned 1 μm apart and stimulated synchronously. The AMPA and NMDA synapse models and parameters were taken from a previous model (Baker et al., 2011; Hyun et al., 2015), fit to experimental results. However, to generate nonlinearity in single dendrite branches the NMDA:AMPA ratio was increased from 0.2 to 2. This higher ratio is common in other models that have investigated NMDA spikes (Poirazi et al., 2003; Larkum et al., 2009; Major et al., 2013). All simulations were run with a timestep of 1 ms.

**Table 2:**
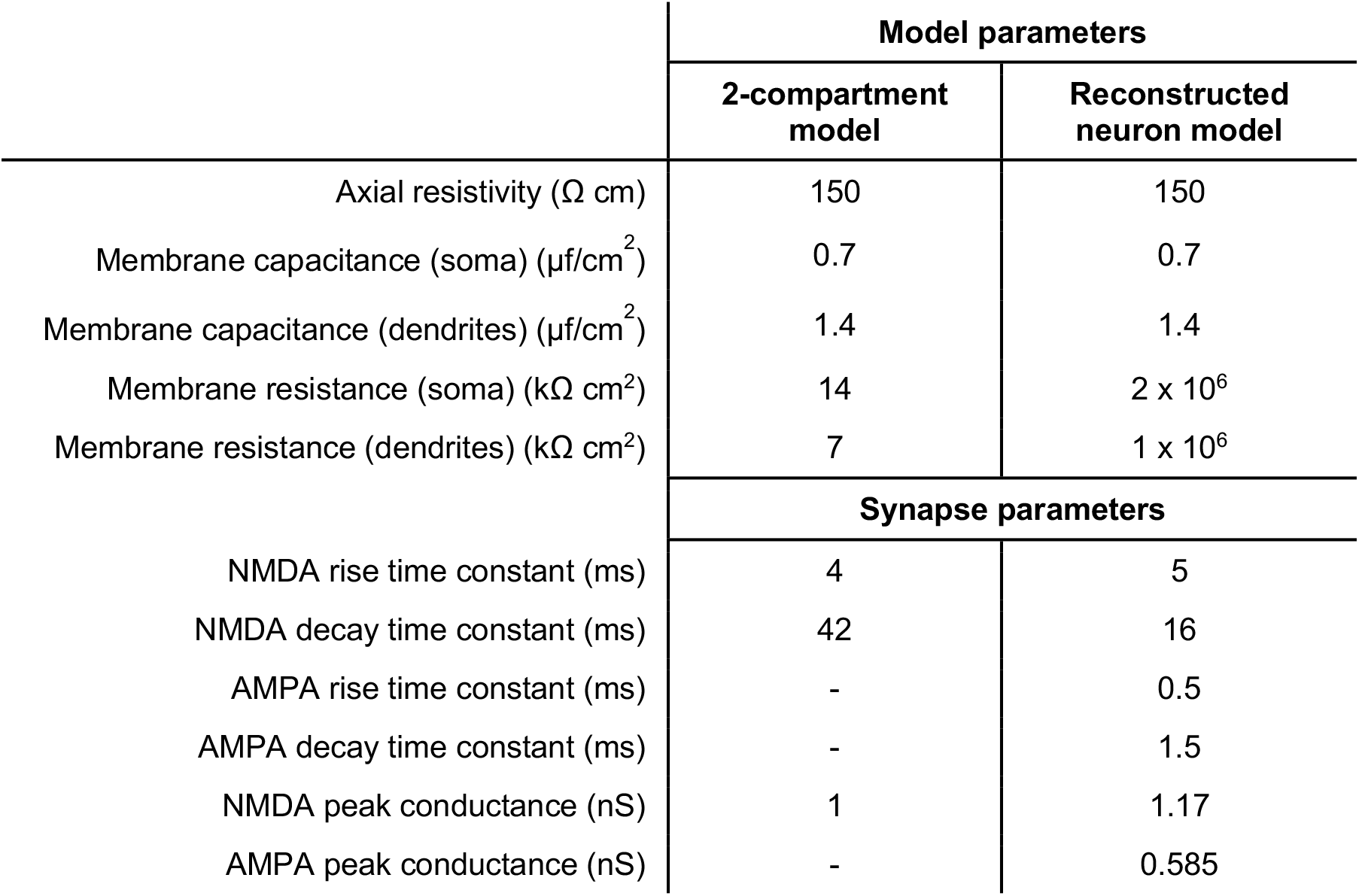
Model and synapse parameters.

### NMDA spike generation and analysis

NMDA spikes were generated by increasing either the synaptic weight of the NMDAR conductance (2-compartment model) or the number of synapses (reconstructed neuron model), from 1 to 20, causing a nonlinear increase in the amplitude of the voltage response. In simulations with AMPA synapses included, we also ran the model with only AMPA synapses (and no NMDA) and subtracted these voltage traces from the original to determine peak the amplitude of the NMDA voltage response. To assess the effect of blocking the potassium channels on the nonlinearity curve, we calculated and compared the maximum slope of the curve and the synaptic weight or number of synapses at the maximum slope.

### Inhibition of potassium channels and simulating acetylcholine

The potassium conductances were blocked in the model by setting their conductance to zero. To simulate acetylcholine, the extent of inhibition for each potassium channel was determined during parameter optimisation. This resulted in a 50% block of the A-type potassium (Ka), M-type potassium (Km), and SK (Kca) conductances and an 80% block of the GIRK (Kir) conductance to simulate acetylcholine.

## Results

### Acetylcholine increases nonlinear synaptic integration in a 2-compartment neuron model

We initially used a 2-compartment model of a soma and dendrite to investigate how acetylcholine may affect NMDAR-mediated nonlinear synaptic integration (Figure 2A). An NMDA-only synapse with peak conductance 1 nS was located on the dendritic compartment and the synaptic weight increased from 1- to 20-fold while recording the resulting voltage in the somatic and dendritic compartments (Figure 2B).

**Figure 2:**
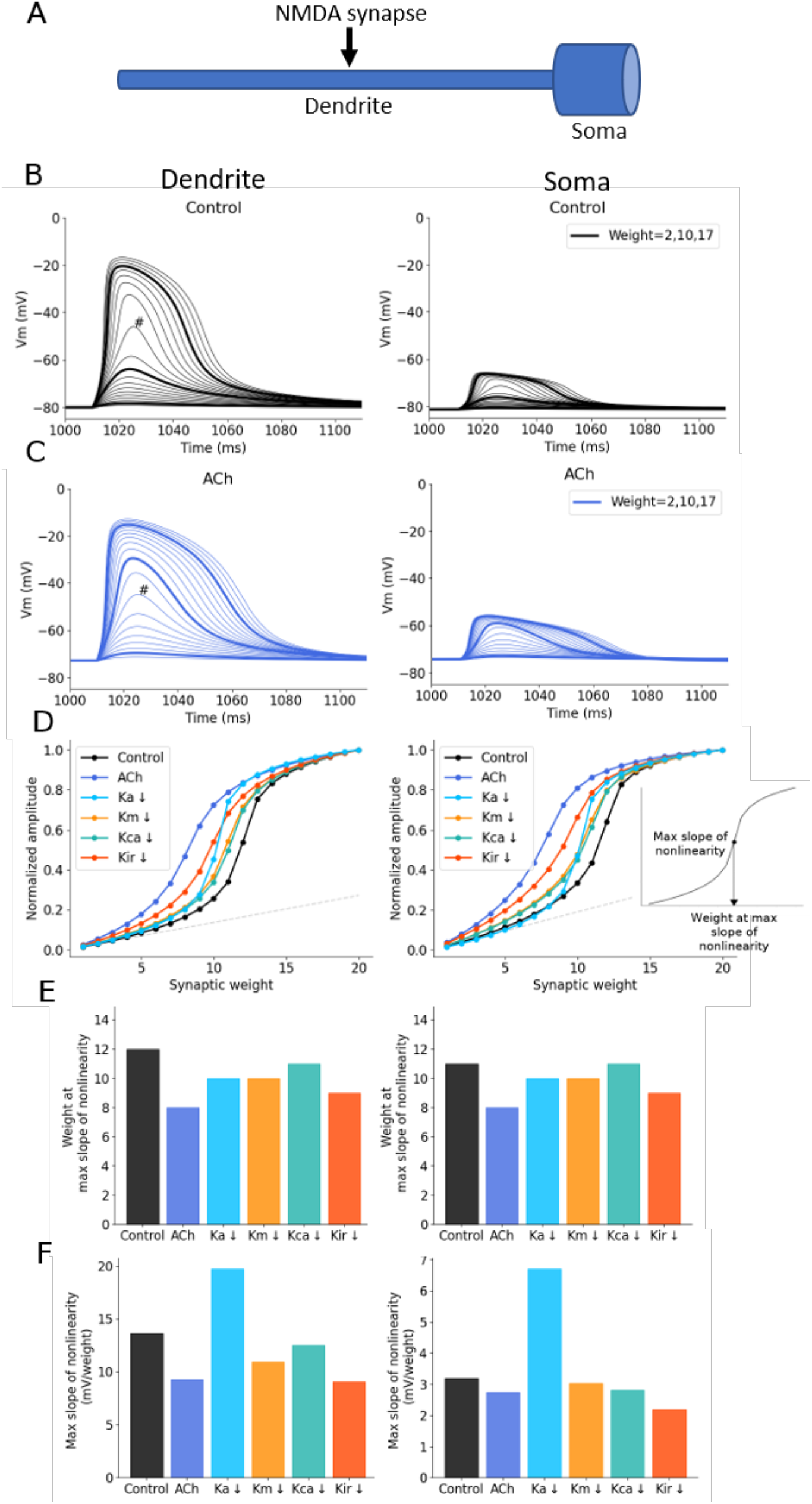
Simulated acetylcholine reduces the synaptic weight at the maximum slope of nonlinearity. (A) Schematic of 2-compartment model. (B) Example voltage traces with increasing synaptic weight measured in the dendritic compartment (left) and somatic compartment (right). # = the weight at the maximum slope of nonlinearity. (C) Example voltage traces with simulated acetylcholine. (D) Effect of inhibiting potassium conductances on the supra-linear increase in voltage response amplitudes with increasing synaptic weight recorded at the dendrite (left) and soma (right). Inset shows how the maximum slope and respective synaptic weight, described in the bar plots in E and F below, are calculated. (E) Change in the synaptic weight at the maximum slope of nonlinearity for each condition. (F) Change in the maximum slope of nonlinearity for each condition.

Increasing synaptic weight caused a nonlinear increase of postsynaptic response (Figure 2D) as expected for voltage-dependent NMDA synapses. We simulated acetylcholine by reducing the potassium conductances Ka, Km, Kca by 50% and Kir by 80%, as determined by parameter optimisation (see Methods). Reduction of these channel conductances in the model increased the resting membrane potential by ~7 mV and the input resistance by ~70% (Figure 2C, D), comparable to experimental data (Figure 1) (Sun and Kapur, 2012). Reducing the potassium conductances increased neuron excitability and enhanced the NMDA response so that less synaptic input was needed to achieve similar amplitude voltage responses in both dendritic and somatic compartments (Figure 2C, D) and shifting the maximum slope of nonlinearity (Figure 2D) from a synaptic weight of 12 to 8 (Figure 2E+F).

To determine individual potassium channel contributions to the increase in nonlinearity seen with acetylcholine, we reduced each potassium channel conductance separately, by 100%. Inhibiting Ka, Km or Kca conductances separately produced a small reduction in the synaptic weight needed to reach nonlinearity, with Kir inhibition causing a slightly larger reduction indicating that no single potassium channel underpinned the effect of acetylcholine (Figure 2D, E). Conversely, inhibition of the A-type potassium conductance caused a considerable increase in the slope of nonlinearity (Figure 2D, F), not observed when blocking any other potassium channel, or when simulating acetylcholine.

### Heterogeneity of nonlinear synaptic integration across dendrites in a reconstructed CA3 neuron model

We next tested the findings from the 2-compartment model in a reconstructed CA3 neuron model. We simulated 1-20 identical synapses, with both AMPA and NMDA components, on individual dendritic branch sections, 1 μm apart, in the SR or SLM region in a reconstructed CA3 pyramidal neuron model (Figure 3A) (see Methods). The synapses on SR dendrites represent recurrent inputs from neighbouring CA3 pyramidal neurons whereas the synapses on SLM dendrites represent inputs from layer II of the entorhinal cortex. The synaptic voltage responses had both a fast AMPAR-based component and a slower NMDAR based component (Figure 3B). To isolate the NMDA component of the voltage response, we calculated the difference in the voltage traces between simulations containing both AMPAR and NMDAR synapses, and simulations containing AMPAR-only synapses. Increasing numbers of synapses on any dendritic branch resulted in monotonically larger voltage responses, increased nonlinear integration, and NMDA spikes (represented by the slower NMDA component of the synaptic response). Notably the threshold for initiation of NMDA spikes varied substantially between dendrites. Overall, the amplitude of the dendritic NMDA response increased with distance of the dendrite section from the soma, whereas the opposite was true for somatic responses, which decreased with dendrite distance from the soma (Figure 3B-E). This was reflected in a shift in the number of synapses required to initiate nonlinear NMDA responses, with fewer synapses generally required to produce nonlinearity in the more distal dendrites.

**Figure 3:**
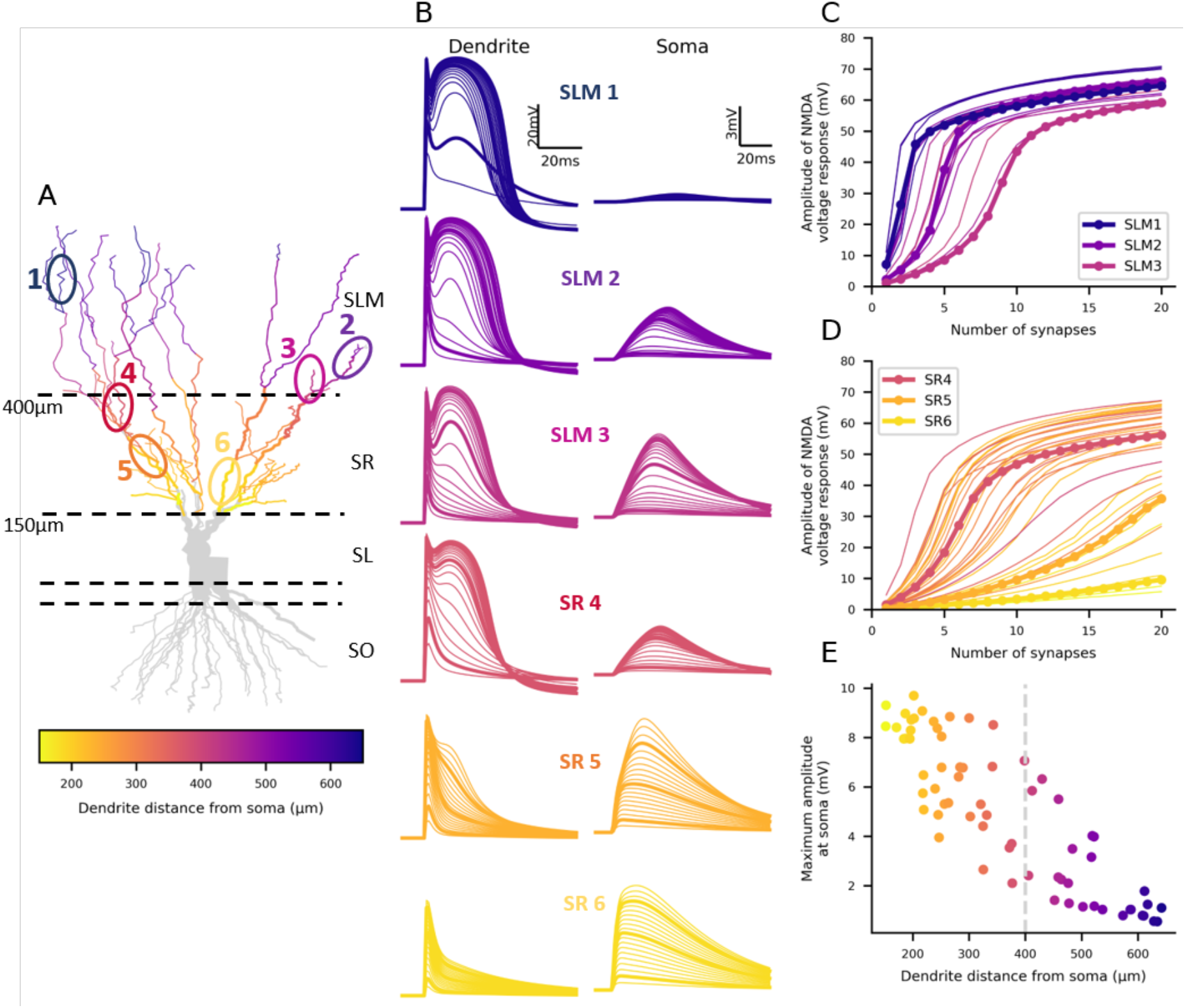
Nonlinearity of synaptic integration varies for dendrites across the proximal-distal axis. (A) Morphology of the reconstructed CA3 neuron model. Dashed lines separate the different strata as labelled on the right with their distance from the soma on the left. Coloured dendrites are the ones included in the generation of the plots in C-E. The circled and numbered sections represent the voltage plot examples in B. (B) Example voltage traces generated by increasing synapse numbers in different dendrite sections recorded in the stimulated dendrite (left) and at the soma (right). Note the different scales for the soma and dendrite recordings. (C) Nonlinearity plots from all the coloured SLM dendrites in A. (D) Nonlinearity plots from all the coloured SR dendrites in A. (E) Maximum voltage response amplitude recorded at the soma decreases with stimulated dendrite distance from the soma.

Subsequently, we investigated the cause of the dendritic variation in nonlinear synaptic integration by regressing the nonlinearity slope and offset against dendritic distance from soma and dendrite local input resistance (Figure 4). The results shown in Figure 3 indicated that increasing dendritic distance from the soma was correlated with fewer synapses at the maximum slope of nonlinearity and we found that this was the case (Figures 4A, B, SR R^2^ = 0.364; SLM R^2^ = 0.496). A possible mechanism for this proximal-distal correlation is increased input resistance in distal dendrites, which would enhance voltage responses to synaptic input and engage nonlinear NMDA conductances. Indeed, the input resistance was well correlated with the number of synapses required for nonlinearity (Figure 4C, SR R^2^ = 0.826; SLM R^2^ = 0.793), showing a tighter relationship than that with dendritic distance. These observed effects were similar for the somatic recordings (Figures 4B, C, right). We also studied the dependence of maximum slope of nonlinearity on distance from the soma (Figures 4D-F). The slope of dendritic nonlinearity increased with distance from the soma (SR R^2^ = 0.250; SLM R^2^ = 0.403) whereas the slope of somatic nonlinearity decreased with distance from the soma, primarily when stimulating the SLM dendrites (Figure 4E, SR R^2^ = 0.018; SLM R^2^ = 0.433) and possibly due to attenuation of the response. Dendritic input resistance was also correlated with the slope of nonlinearity (Figure 4F, SR R^2^ = 0.883; SLM R^2^ = 0.587). To summarise, dendritic distance from the soma and input resistance can explain the differences in the generation of NMDA nonlinearity between dendrites.

**Figure 4:**
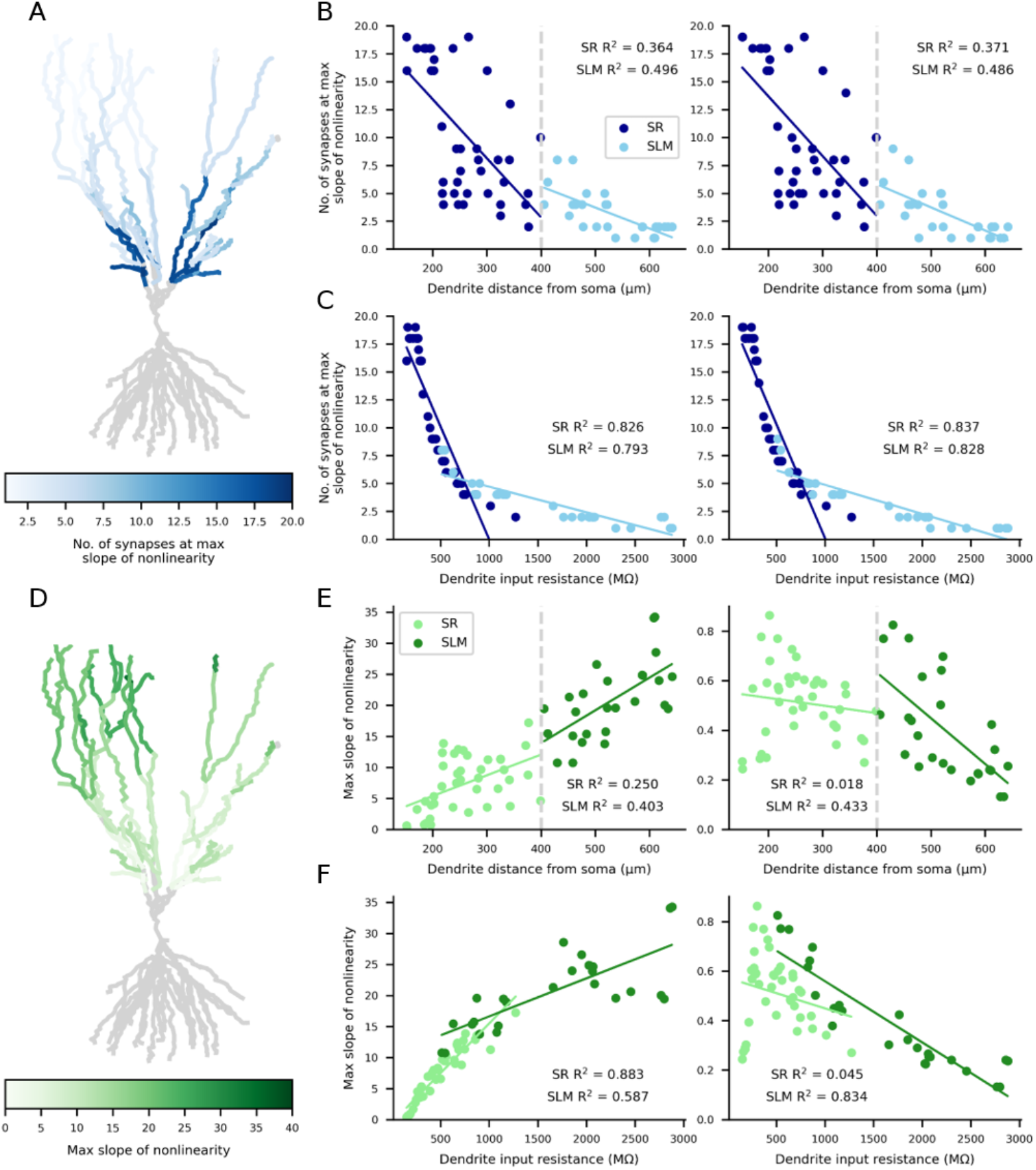
The number of synapses at the maximum slope of nonlinearity reduces with distance from the soma due to increased dendritic input resistance. (A+B) Number of synapses at the maximum slope of nonlinearity decreases with dendrite distance from the soma. Colour coding for entire neuron shown in A with individual dendrites quantified in B. (C) An increasing dendritic input resistance reduces the number of synapses at the maximum slope of nonlinearity. (D+E) The maximum slope of nonlinearity increases with distance from the soma as recorded in the dendrite, however when recorded at the soma (E, right), the maximum slope decreases in the SLM dendrites with distance from the soma. Colour coding for entire neuron shown in D with individual dendrites quantified in E. (F) An increasing dendritic input resistance increases the maximum slope of nonlinearity in the dendrite (left) but reduces the maximum slope in the soma (right).

### Acetylcholine boosts nonlinear synaptic integration in the reconstructed neuron model

Next, we investigated the impact of acetylcholine on nonlinear synaptic integration in the reconstructed neuron (Figure 5). We simulated acetylcholine as described in the 2-compartment model. Similar to the 2-compartment model, acetylcholine depolarised the neuron’s resting potential and enhanced voltage responses to synaptic stimulation (Figure 5B). This was reflected in a reduction in the number of synapses required to generate nonlinear synaptic integration in each of the example dendrites (Figure 5C). The magnitude of the effect varied across dendrites but was true for the majority of both SR and SLM dendrites, with up to 5 fewer synapses needed for nonlinearity in the SR dendrites, and up to 2 fewer synapses in the SLM dendrites (Figure 5D). Furthermore, acetylcholine reduced the slope of nonlinearity in most dendrites, similar to the 2-compartment model (Figure 5E).

**Figure 5:**
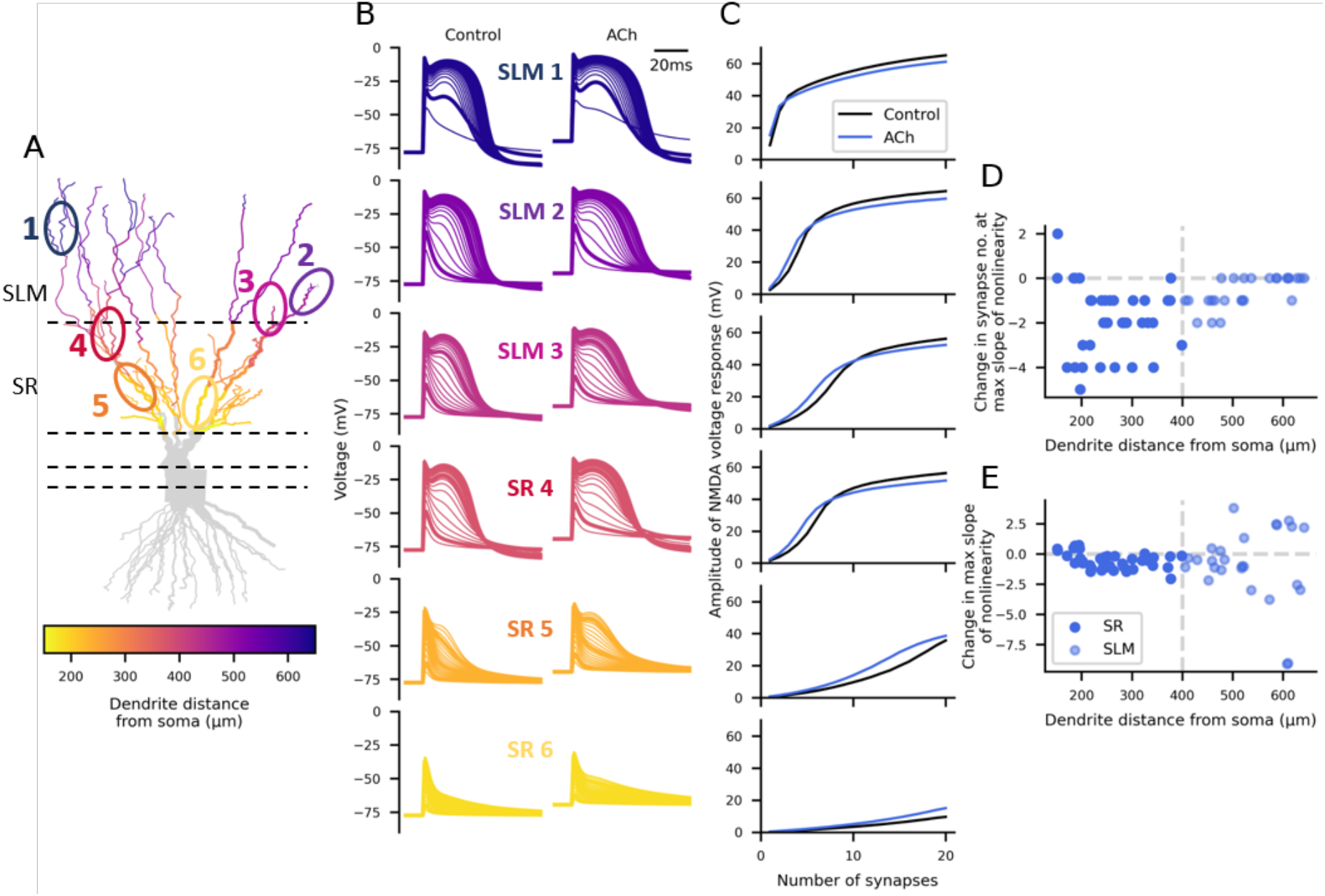
Acetylcholine boosts nonlinear synaptic integration in most dendrites. (A) Morphology of the reconstructed CA3 neuron model. Coloured dendrites are the ones included in the generation of the plots in D and E. The circled and numbered sections represent the voltage plot examples and nonlinear curves in B and C. (B) Example dendritic voltage traces of increasing synapse numbers in different dendrite sections recorded in the stimulated dendrite (left) and then with simulated acetylcholine (right). (C) Nonlinearity curves for control (black) and acetylcholine (blue) for each example dendrite in B. (D) The change in number of synapses at the maximum slope of nonlinearity with acetylcholine for all SR and SLM dendrites. (E) The change in the maximum slope of nonlinearity with acetylcholine for all the SR and SLM dendrites.

We then calculated the average effect of acetylcholine across all SR and SLM dendrites, as well as the inhibition of individual potassium channel conductances (Figure 6). On average, acetylcholine reduced the number of synapses required to initiate nonlinearity from 2.50 ± 0.37 to 1.88 ± 0.25 in the SLM dendrites and from 10.30 ± 0.92 to 8.50 ± 0.88 in the SR dendrites (mean ± standard error; Figure 6A). This effect was similar in the somatic response (SLM: from 2.50 ± 0.38 to 1.88 ± 0.25; SR: from 10.53 ± 0.92 to 8.70 ± 0.88). Again, no one particular potassium conductance could fully account for the effect of acetylcholine. In the 2-compartment model, blocking the Ka channel increased the slope of nonlinearity. We also found this effect in the reconstructed neuron model, to a much lesser extent, and only in the SLM dendrites (Figure 6B). This is likely because Ka channel density increases with distance from the soma in the model in line with experimental data (Kim et al., 2012). Therefore, inhibition of Ka conductances in the distal SLM dendrites causes a greater effect. Blocking the Kca channel also seemed to slightly increase the slope of nonlinearity, but at the soma, as opposed to in the dendrites. To summarise, the results in the reconstructed neuron model reflect the conclusions from the 2-compartment model, in which acetylcholine reduces the number of synapses required for nonlinearity and blocking the Ka conductance increases the slope of nonlinearity.

**Figure 6:**
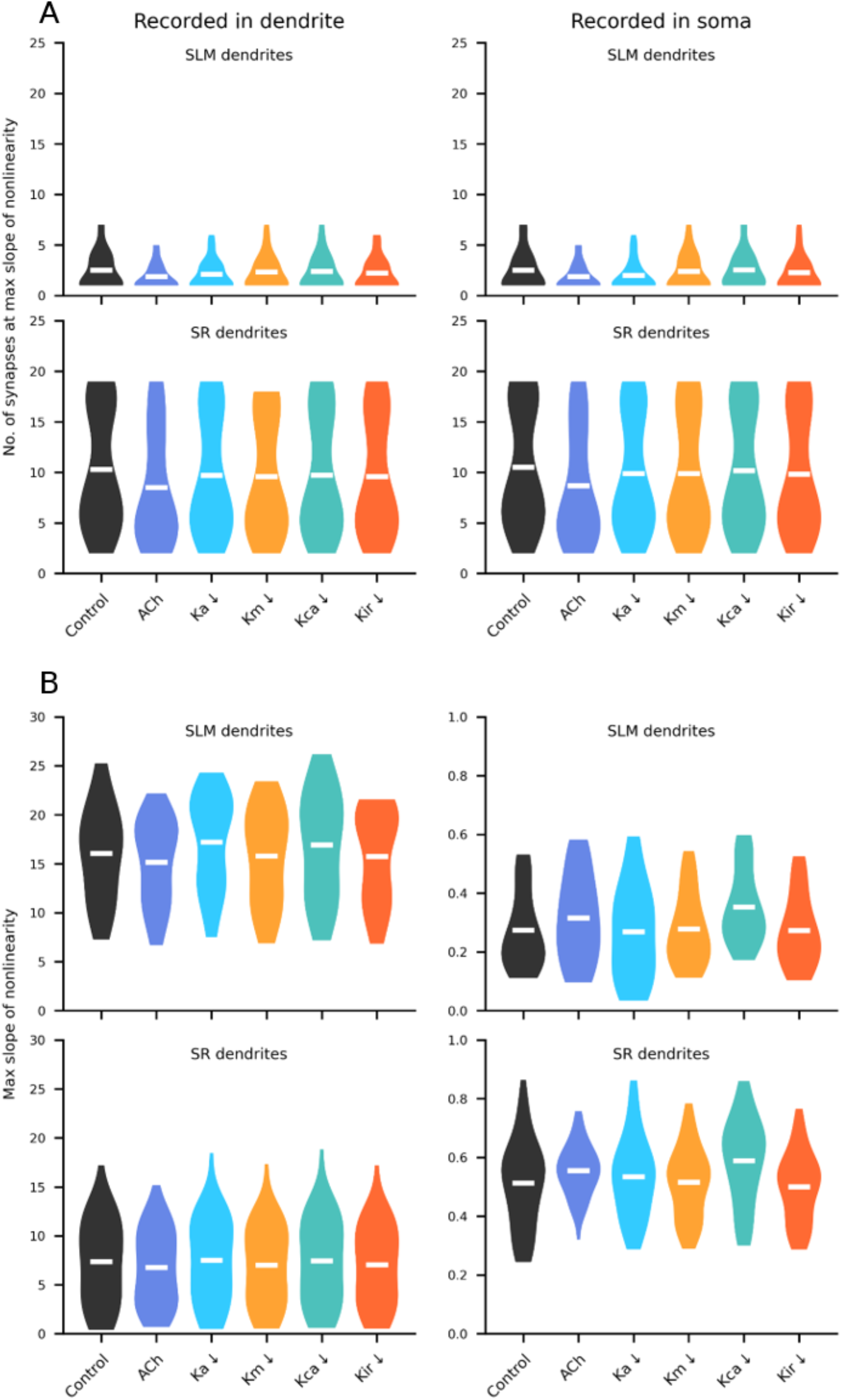
Acetylcholine reduces the number of synapses at the maximum slope of nonlinearity in the reconstructed neuron model. Violin plots of density of measurements across all dendrites, white bars represent the mean. (A) Acetylcholine reduces the number of synapses at the maximum slope of nonlinearity in SLM (top) and SR (bottom) dendrites. (B) Change in the maximum slope of nonlinearity for each condition in the SLM (top) and SR

## Discussion

In a reconstructed CA3 neuron model, we have shown that simulating acetylcholine reduced the number of synapses required for nonlinear integration of glutamatergic synaptic inputs in SR and SLM dendrites. We have also demonstrated that dendrite distance from the soma and input resistance correlate with the dendrite’s ability to generate nonlinearity, with distal dendrites with a higher input resistance more readily producing nonlinearity. Additionally, the A-type potassium channel (Ka) modulated the slope of nonlinearity in the distal SLM dendrites, where there is a higher proportion of Ka channels.

Nonlinear synaptic integration has previously been studied in CA3 pyramidal neurons *ex vivo*. Makara and Magee (2013) observed NMDAR-mediated nonlinearity in SR and SO CA3 dendrites with glutamate uncaging. In terms of variability between dendrite branches, they characterized two populations with a fast or slow NMDA spike decay time. They did not report a correlation between dendrite distance from the soma and NMDA spike decay time but determined the GIRK channel as primarily responsible for regulating the decay time. Here, we did not analyse the NMDA spike decay time but there did not seem to be a major effect from blocking the potassium channels individually on the onset of nonlinearity. However, of all the channels, blocking the Kir channel displayed the greatest impact on the nonlinearity threshold and therefore could play an important role in the modulation of NMDA spikes. Kim et al., (2012) analysed dendritic sodium spikes in CA3 neurons and reported a reduced threshold with increasing distance from the soma, in accordance with the NMDA spike threshold results displayed here. Differences in dendrite ability to generate sodium spikes has also been noted in CA1 oblique dendrites (Losonczy et al., 2008) where distinct strong or weak branch populations were identified. However, in contrast with the data from CA3 neurons, the branches closer to the soma were more readily able to generate dendritic sodium spikes. This variation in dendrite ability to generate dendritic spikes suggests that certain synaptic inputs can be prioritised.

The effect of acetylcholine on dendritic integration has been previously investigated in the somatosensory cortex (Williams and Fletcher, 2019). Optogenetically released acetylcholine increased apical dendrite excitability, facilitated the integration of inputs, and caused long duration dendritic plateau potentials that impacted action potential burst firing. This cholinergic modulation of dendritic integration was mediated by the facilitation of R-type Ca^2+^ channels. The cholinergic enhancement of dendritic excitability and integration supports the results shown here. Furthermore, in CA1 neurons (Losonczy et al., 2008), 5 μM carbachol, combined with repetitive spike generation in weak branches, selectively enhanced the weak branch sodium spike strength and its propagation into stronger proximal branches. This is in line with the data shown here in which acetylcholine affects the SR dendrites, with a higher NMDA spike threshold, more so than the SLM dendrites that are already capable of generating nonlinearity with minimal synaptic input. This suggests that acetylcholine selectively facilitates NMDA spike generation in the SR dendrites and could therefore impact CA3-CA3 recurrent synaptic plasticity by increasing Ca^2+^ influx to synapses from both local voltage-gated Ca^2+^ channels and from the NMDARs themselves. Acetylcholine also alters somatic excitability of CA3 pyramidal neurons but the nature of the modulation depends on the type of pyramidal neuron. Athorny, burst firing neurons reduce their burst firing in response to acetylcholine whereas thorny regular firing neurons increase their firing rate (Hunt et al., 2018). This may reflect varying balances of ion channel conductances at the soma and it is not clear if these are also found in the dendrites.

We also found that the A-type potassium channel regulated the slope of nonlinearity. This measure does not relate to the absolute number of synapses needed to reach the nonlinearity, but rather the steepness of the response amplitude as further synapses are activated. This result reflects a previous study in which blocking the A-type potassium channel increased the dendritic plateau potential amplitude in CA1 apical dendrites (Cai et al., 2004). Downregulation of this channel also increased the weak branch sodium spike strength in CA1 dendrites (Losonczy et al., 2008), highlighting its importance in CA1 nonlinear integration.

### Functional implications

This study predicts that acetylcholine facilitates NMDA spike generation in CA3 dendrites. NMDA spikes have been shown to be necessary for generating timing-dependent associative plasticity at CA3-CA3 recurrent synapses (Brandalise et al., 2016). Therefore, the presence of acetylcholine could facilitate this plasticity, by lowering the threshold for NMDA spike initiation, and strengthen these connections to enable new memory ensembles to form. Alternatively, promotion of spiking in CA3 neurons by NMDA spikes (Makara and Magee, 2013) could induce spike timing-dependent plasticity at CA3-CA3 recurrent synapses (Mishra et al., 2016).

NMDAR-dependent dendritic plateau potentials that mediate nonlinear integration have been shown to induce plasticity that underlies CA1 place field firing (Bittner et al., 2015, 2017). Neuromodulatory input form the locus coeruleus has been shown to be important for promoting CA1 place cell formation around new reward locations (Kaufman et al., 2020), as well as enhancing CA1 dendritic excitability and LTP (Liu et al., 2017; Bacon et al., 2020). Taking into consideration these studies and the results shown here, acetylcholine could also be implicated in place cell generation by facilitating dendritic excitability and NMDAR-mediated plateau potentials (Prince et al., 2016; Fernández de Sevilla et al., 2020).

Interestingly, acetylcholine had a heterogeneous effect on excitability of different dendrites. In some dendrites it lowered the NMDA spike threshold by ~50%, while in others it caused no reduction in threshold. This heterogeneity implies that acetylcholine release could shift the relative sensitivities of somatic voltage to synaptic inputs on different dendrites, as opposed to just implementing a uniform increase in sensitivity to all inputs. Future studies on the effects of acetylcholine in neural circuits could address how this effect would modulate information flow and ensemble formation in CA3 networks.

In conclusion, this study predicts that acetylcholine facilitates the generation of NMDA spikes in CA3 pyramidal neuron dendrites, which could provide an important mechanism for the storage of new memories in hippocampal circuitry.

## Acknowledgements

We thank Simonas Griesius and Rahul Gupta for helpful discussions. This work was supported by Wellcome Trust (RH PhD studentship, JRM 101029/Z/13/Z), BBSRC (JRM, BB/R002177/1), MRC (COD, MR/S026630/1), and Leverhulme Trust (COD, RPG-2019-229).

